# Dense geographic and genomic sampling reveals paraphyly and a cryptic lineage in a classic sibling species complex

**DOI:** 10.1101/491688

**Authors:** Ethan Linck, Kevin Epperly, Paul van Els, Garth M. Spellman, Robert W. Bryson, John E. McCormack, Ricardo Canales-del-Castillo, John Klicka

## Abstract

Incomplete or geographically biased sampling poses significant problems for research in phylogeography, population genetics, phylogenetics, and species delimitation. Despite the power of using genome-wide genetic markers in systematics and related fields, approaches such as the multispecies coalescent remain unable to easily account for unsampled lineages. The *Empidonax difficilis* / *E. occidentalis* complex of small tyrannid flycatchers (Aves: Tyrannidae) is a classic example of a widely distributed species with limited phenotypic geographic variation that was broken into two largely cryptic (or “sibling”) lineages following extensive study. Though the group is well-characterized north of the U.S. Mexico border, the evolutionary distinctiveness and phylogenetic relationships of southern populations remain obscure. In this paper, we use dense genomic and geographic sampling across the majority of the range of the *E. difficilis* / *E. occidentalis* complex to assess whether current taxonomy and species limits reflect underlying evolutionary patterns, or whether they are an artifact of historically biased or incomplete sampling. We find that additional samples from Mexico render the widely recognized species-level lineage *E. occidentalis* paraphyletic, though it retains support in the best-fit species delimitation model from clustering analyses. We further identify a highly divergent unrecognized lineage in a previously unsampled portion of the group’s range, which a cline analysis suggests is more reproductively isolated than the currently recognized species *E. difficilis* and *E. occidentalis.* Our phylogeny supports a southern origin of these taxa. Our results highlight the pervasive impacts of biased geographic sampling, even in well-studied vertebrate groups like birds, and illustrate what is a common problem when attempting to define species in the face of recent divergence and reticulate evolution.

## Introduction

The effects of incomplete or biased taxon sampling are a perennial concern in evolutionary biology (Hillis 1998, Heath et al. 2008; Rosenberg et al. 2005; Schwartz and McKelvey 2009; Puechmaille 2016). In phylogenetics, missing taxa can pose significant problems for tree estimation (Felsenstein 1978; Hillis 1998; Reddy et al. 2017), with cascading consequences for macroevolutionary analysis, biogeography and studies of trait evolution (e.g. FitzJohn et al. 2009, Ahrens et al. 2016). In population genetics, sampling scheme can drive spurious clustering and influence the value of summary statistics (Rosenberg et al. 2005; Schwartz and McKelvery 2009; Chikhi et al. 2010), in turn biasing inference of selection (Excoffier et al. 2009) and genome-wide association studies (‘GWAS’; Astle and Balding 2009).

For systematists studying nonmodel organisms, this source of error is often compounded by geographic biases in biological knowledge. Both range size and body size can be correlated with the probability of a given species being described (Blackburn and Gaston 1995; Collen et al. 2004), but are also variables with strong relationships to latitude in many taxa (e.g. Stevens 1989; Freckleton et al. 2003; Riemer et al. 2018). Taxonomic practices and species concepts also vary across space: species are often more narrowly defined (or “split”) at higher latitudes (Jones et al. 2012, Barrowclough et al. 2016), in part reflecting the higher density of scientific institutions in Europe and North America.

As species are widely cited as the fundamental unit of biodiversity (Hull 1977, Coyne 1994, Magurran 2013, Hohenegger 2014; de Queiroz 2005), the implications of misdiagnosis or unevenly applied diagnostic criteria are significant. When logistical or political constraints prevent wider geographic sampling, taxonomic decisions are forced to rely on samples from an incomplete or unrepresentative portion of a given species’ range (Avendaño et al. 2017). Given that lower latitudes are likely to contain greater genetic diversity and a greater number of distinct lineages (Harvey and Brumfield 2015, Smith et al. 2014, Barrowclough et al. 2016, Smith et al. 2017), species with large ranges that were originally delimited from sampling biased towards higher latitudes may eventually prove paraphyletic (McKay and Zink 2010; Zucker et al. 2016), requiring revised taxonomic treatments that affect biodiversity estimates and conservation priorities along with the numerous disciplines dependent on accurate species lists (Jones et al. 2012, Barrowclough 2016).

These concerns are especially salient for delimitation of cryptic and sibling species, where the absence of easily diagnosable morphological variation across ranges limits the ability to make informed choices about the potential significance of sample exclusion on inference. Molecular phylogenetics has revolutionized the study of cryptic and sibling lineages (e.g. Knowlton 1993; de Leon and Nadler 2010; Adams et al. 2014; Papakostas et al. 2016) and substantially altered our understanding of the relationship between reproductive isolation and phenotype, geography, and genetic variation (Bickford et al. 2007; Fišer et al. 2018). The recent emergence of model-based species delimitation with genome-wide genetic markers aids in reducing bias due to arbitrary divergence thresholds and the stochasticity of a small sample of genes (Leaché et al. 2014; Hotaling et al. 2016). However, these approaches remain unable to correct for phylogenetic relationships with unsampled lineages, underscoring the continued importance of geography to systematic biology.

Tyrannid flycatchers (Aves: Tyrannidae) are the largest bird family in the New World, and have long posed problems for taxonomists and systematists due to limited morphological variation in many groups, conflicting characters, and evidence of convergent evolution (Traylor 1977; Birdsley 2002; Chaves et al. 2008). Of the tyrannids, the *Empidonax difficilis* / *E. occidentalis* complex provides a representative and classic example of widely-distributed species with limited geographic variation, broken into separate, largely cryptic, lineages only after extensive study (Johnson 1980, Zink and Johnson 1984, Johnson and Martin 1988). Based on quantitative analysis of museum specimens and vocalizations across its full range, in addition to putative assortative mating at a contact zone, the Western Flycatcher *E. difficilis* was split into interior western North American (Cordilleran Flycatcher, *E. occidentalis)* and coastal western North American (Pacific Flycatcher, *E. difficilis)* species (Johnson 1980), themselves sister to the allopatric species *E. flavescens* (Yellowish Flycatcher) which occurs throughout Central American highlands, largely east of the Isthmus of

Tehuantepec. Subsequent population genetic analysis using allozyme data supported the reciprocal monophyly of these groups, though sampling was restricted to populations in the United States (Johnson and Marten 1988). Both an analysis of a contact zone in interior southwest Canada (Rush et al. 2009) and unpublished mtDNA and nuDNA phylogeography studies (Klicka et al. unpublished; Rush et al. in prep) suggest locally extensive hybridization between forms, but to date, the affinities of resident and migratory populations in Mexico remain unexamined with genetic data.

Here, we explore the influence of biased sampling on determining species limits in this complex of *Empidonax* flycatchers. We combine dense population sampling with RADseq, UCE, and mtDNA sequence data from across the majority of the latitudinal range of the currently recognized species *E. difficilis* and *E. occidentalis.* Using these data, we ask whether comprehensive geographic and genomic sampling changes estimated phylogenetic relationships and species limits in a classic sibling species complex, shedding light on our ability to infer fundamental evolutionary units and processes in morphologically conserved, geographically widespread taxa.

## Materials and Methods

### Sampling, Library Preparation, and Sequencing

We collected 308 vouchered tissues from four *Empidonax species*, with localities in Canada, the United States, Mexico, Guatemala, Honduras, and Panama (**Figure 1; Table S1**). Of these, 295 belonged to our ingroup, the *E. difficilis* / *E. occidentalis* complex. We represented the entirety of the group’s geographic range with the exception of *E. difficilis* populations in northern British Columbia and Alaska and *E. occidentalis* populations in southeastern British Columbia and Alberta, a region previously examined in detail by Rush et al. 2009. The remaining 13 samples in our study belonged to outgroup taxa: *E. flaviventris* (n = 4), a long-distance migrant that breeds in North American boreal habitats, and *E. flavescens* (n = 9), a Central American endemic.

**Figure 1:**
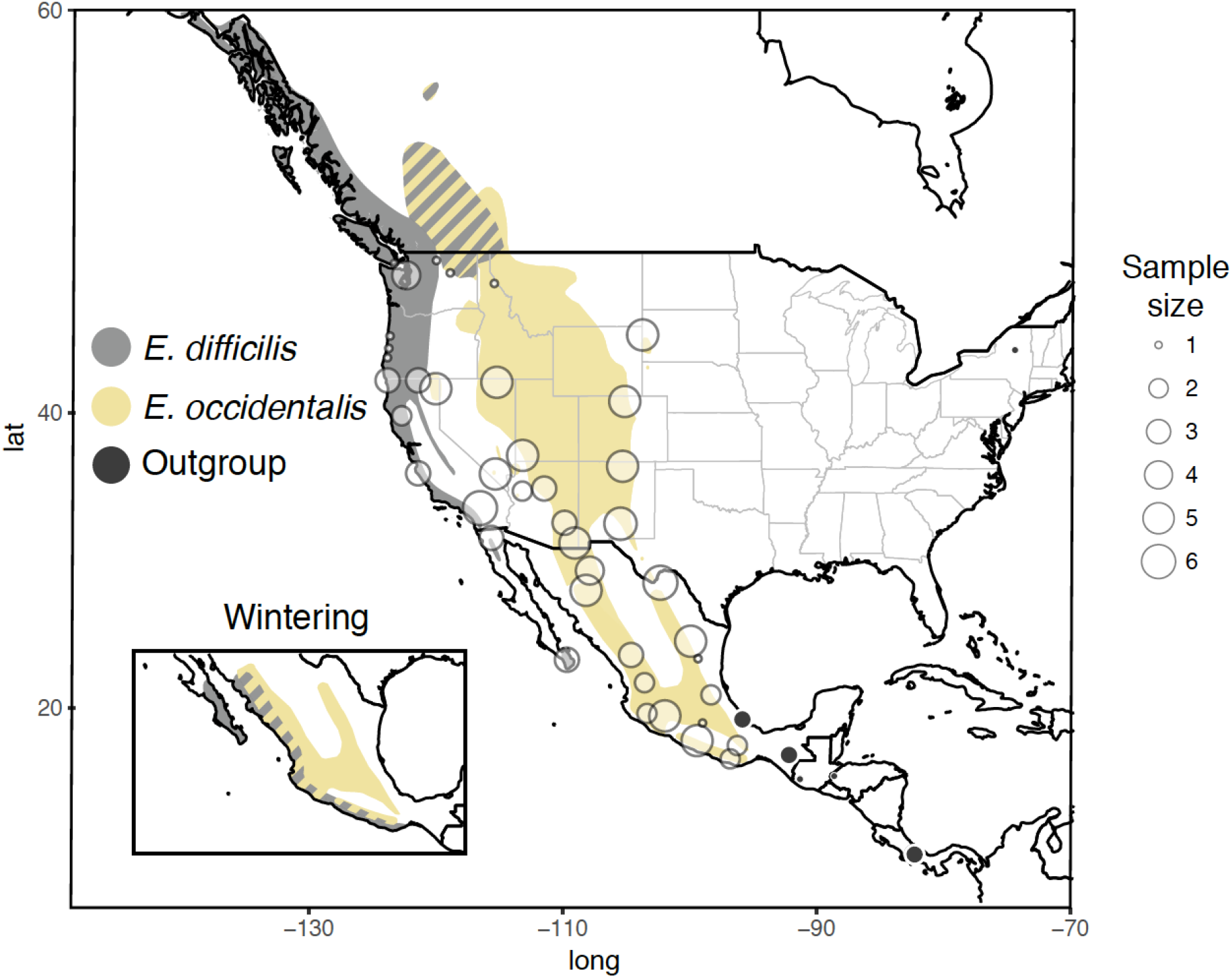
Localities and sample size for the *Empidonax difficilis* / *E. occidentalis* complex, with outgroups *E. flaviventris* and *E. flavescens.*

We extracted total genomic DNA from all samples using a Qiagen DNeasy tissue extraction kit, following the manufacturer’s recommended protocol. For all samples, we sequenced 1041 bp of the mitochondrial gene NADH dehydrogenase subunit 2 (ND2), using primers L5215 (Hackett 1996) and TrC (Miller et al. 2007) and 12.5 *μL* PCR reactions on a T100 thermal cycler (Bio-Rad, Hercules, CA) as follows: 94°C for 2.5 min, 35 cycles of (94°C for 30 s, 54°C for 30 s, 72°C for 1 min), 10 min at 72°C, 10°C hold. We sent PCR products to the High-Throughput Genomics Unit at the University of Washington for cleanup and sequencing.

Based on fragment size and DNA concentration, we then chose a 144-sample subset for reduced representation genome sequencing with the double digest restriction-associated DNA sequencing library preparation method (Peterson et al. 2012). This subset included 134 ingroup samples (46 *E. difficilis,* 88 *E. occidentalis)* and 10 outgroup samples (two *E. flaviventris,* eight *E. flavescens)* representing all four currently recognized species mentioned above. We used digestion enzymes `Sbf1` and `Msp1` and a Pippin Prep size selection device to retain fragments between 415 and 515 bp. Our libraries were then sequenced for 100 bp single-end reads on an Illumina HiSeq 2500 at University of California Berkeley’s QB3 Vincent J. Coates Genomics Sequencing Laboratory. After sequencing, we excluded three ingroup samples and one outgroup sample *(E. flaviventris)* due to poor data quality.

Finally, we chose a second, nonexclusive subset of 37 individuals for the reduced representation sequencing approach of targeted capture for ultraconserved elements and their flanking regions (UCEs; Smith et al. 2014). This subset focused on the southwestern US, Mexico, and Central America. It also included all four species, with 34 ingroup samples (1 *E. difficilis,* 33 *E. occidentalis)* and four outgroup samples (1 *E. flaviventris,* 3 *E. flavescens).* We sent raw extractions to RAPiD Genomics (Gainesville, FL USA), where they enriched each pool using a set of 5,472 custom-designed probes (MYbaits; MYcroarray, Inc., Ann Arbor, MI, USA) targeting 5,060 UCE loci (Faircloth et al. 2012; a full protocol can be found on http://ultraconserved.org.) Pooled libraries were then sent to the University of Florida ICBR Facility for 150 bp read paired-end sequencing on an Illumina NextSeq500.

### Sequence Assembly

We edited and aligned forward and reverse chromatograms from ND2 sequences using `Sequencher` v. 4.9 and the `MUSCLE` alignment algorithm implemented in `MEGA` v. 7.0 (Kumar, Stecher, and Tamura 2015). We performed *de novo* sequence assembly and variant detection from ddRADseq reads using ipyrad v. 0.7.8 (Eaton 2014), implementing a clustering threshold of 85% and a minimum read depth of 6. To test the effect of increased phylogenetic distance on SNP number and alignment completeness, we repeated all steps in the `ipyrad` pipeline with outgroup individuals excluded, ultimately performing all phylogenetic analyses with the full set of taxa, and population assignment analyses using only the ingroup. Our final sequence alignment only included loci where no more than 50% of individuals were missing, and only included individuals with more than 50% of all loci sequenced, resulting in the loss of three samples. To exclude putative paralogs, we dropped loci with more than 5 heterozygous sites, and included no loci sharing a heterozygous site across more than 50% of individuals. We assembled UCE reads using a modified version of the `PHYLUCE` pipeline (Faircloth 2015), described in detail elsewhere (Bryson et al. 2017), and called SNPs using the custom pipeline described in Zarza et al. (2018), with our most completely assembled sample (UWBM 118082; *E. occidentalis* from Nuevo León, MX) serving as the reference.

### Phylogenetic Inference

For each of our three datasets, we inferred phylogenetic relationships among taxa with two methods. We first inferred maximum-likelihood phylogenies in `RAxML` v 8.2.4 (Stamatakis 2006) via the `CIPRES` portal (Miller et al. 2010). After determining the best-fit model of sequence evolution for our concatenated data matrices (`GTRGAMMA`), we ran the program for 1000 bootstrap replicates to estimate node support.

We next used `SVDquartets` (Chifman and Kubatko 2014), implemented in `PAUP` v 4.0 (Swofford 2002), to estimate both a phylogeny of individual lineages and a species tree under a coalescent model robust to the influence of gene flow between sister lineages (Long and Kubatko 2018). `SVDquartets` randomly samples quartets (sets of four taxa, or tips) from the data matrix and calculates the singular value decomposition score to evaluate their optimal relationship. The program then combines these quartets algebraically to infer the overall species tree, and estimates branch support with 100 nonparametric bootstrap replicates. For our ddRADseq data, we additionally inferred a species tree using only individuals determined to represent “pure” populations, which we defined as ≤ 5% admixed, based on our `structure` analysis (see below).

### Population Structure Inference

To identify genetically differentiated populations (or species) and identify putative regions of admixture among them, we performed multivariate ordination and Bayesian coalescent clustering on ddRADseq and UCE SNP datasets independently. We conducted a principal-components analysis (PCA; Pearson 1901) using the covariance matrix of allele frequencies in each sample. We then identified putative genetic clusters with *k*-means clustering using the `R` package `adegenet` v2.0.1 (Jombart 2008), setting the maximum number of clusters to 10 and retaining all principal components. Next, we conducted Bayesian clustering under a coalescent model allowing for admixture using the program `structure` (Pritchard et al. 2000). Because including singletons can strongly bias inference of population structure (Linck and Battey 2018), we used a custom script to restrict our data matrix to only parsimony-informative SNPs (https://github.com/slager/structure_parsimony_informative). We employed default priors, correlated allele frequencies, and stipulated a chain length of 1,000,000 with the first 100,000 steps discarded as burn-in. We replicated structure analyses 10 times for each value of *k* from 1-8, assessed change in marginal likelihood across values of *k* using `STRUCTURE HARVESTER` (Earl 2012), and used `CLUMPP` (Jakobsson and Rosenberg 2007) to average permuted matrices across replicates while controlling for label switching. We plotted individual ancestry proportions using `structurePlotter` v 1.0 (Battey 2017).

Because `structure` may not reliably infer population structure when population sample sizes are uneven (Puechmaille 2016), we also performed all population structure analyses with replicated subsampling. We first assigned samples to putative populations based on their assigned *k*-means cluster, using the subdivision scenario where Bayesian information criterion (BIC) showed the greatest change in value (*k*=4 for both datasets). From each of these populations, we next randomly selected the number individuals equal to sampling the smallest population (n=7 for ddRADseq; n=2 for UCEs), while retaining admixed individuals. For the RADseq dataset, we repeated this process until all individuals were included in at least two of the subsets resulting in a total of 18 subsets. These data matrices were then used for PCA / *k*-means clustering and `structure` analyses with identical parameters to those described above, with the final RADseq `structure` results plotted from the averaged *Q*-value for each individual from across all subset runs.

### Genomic Cline Analysis

To assess the relative level of reproductive isolation between pairs of taxa with adjacent geographic ranges, we analyzed the width and relative position of genomic clines using the `R` package `HZAR` (Derryberry et al. 2013). We selected three transects across regions identified as hybrid zones in our population structure analyses, representing contact points between USE and USW, USW and MXO, and MXO and MXS (**Figure S6**; see **Results** section for definitions of regional abbreviations).

We first calculated total transect length and the distance between each transect locality using the `R` package `geosphere` (Hijmans 2017). Using averaged *Q*-value from structure analyses using our 18 individual subsampled matrix, we selected scores for each species as follows: for Transect 1 (between USW and USE), we used *Q*-values from USE; for Transect 2 (between USE and MXO), we used *Q*-values from MXO; and for Transect 3 (between MXO and MXS), we used *Q*-values from MXS. Setting maximum cline width to +/-100 km of geographic distance sampled to increase the efficiency of the algorithm, we ran 10 models. These consisted of the presence of exponential decay on both ends of the cline in isolation, on both ends of the cline together, or the absence of exponential decay on either end of the cline, repeated with a fixed and with a free scaling parameter.

Checking to ensure the convergence of each chain, we ran 3 MCMC chains for 1,000,000 generations with 10,000 generations of burn-in for each model. We then concatenated results from each independent chain for each model, and selected among our competing models using corrected Akaike Information Criterion (AICc) scores. We extracted maximum likelihood cline center and width values from the best model, expressing confidence intervals of each parameter as 2 log-likelihoods. We modeled maximum likelihood clines using 95% of the credible cline region.

## Results

### Sequencing and Assembly

We successfully amplified and sequenced ND2 for all 308 samples. Illumina sequencing of our ddRADseq libraries returned an average of 1,241,201 quality filtered reads per sample. Clustering within samples while including outgroup taxa resulted in a total of 594,004 putative loci, with an average depth of coverage of 26.7. After clustering across individuals and applying paralog and depth-of-coverage filters, we retained an average of 6,880 loci per sample, which were 631,934 bp long when concatenated. Variant detection resulted in a total of 50,210 SNPs, which we reduced to 6,808 unlinked SNPs. After applying a filter to select only parsimony-informative sites, this number was further reduced to 3,427 SNPs. Excluding outgroup taxa, we identified a total of 55,410 putative loci with an average depth of coverage of 27.2. Clustering across individuals and filtering reduced this to 7,319 loci and 670,994 bp. These loci contained 44,757 total SNPs, 7,194 unlinked SNPs, and 4,048 unlinked parsimony informative SNPs. Our four evenly subsampled alignments ranged from 1,864 to 1,910 parsimony informative SNPs.

UCE libraries returned an average of 1,404,000 quality filtered reads per sample. After assembly and alignment to probe sequences, our data matrix included 4,203 UCE loci from all samples for a total concatenated length of 2,066,792 bp. Mean locus length was 470 bp, with a range of 162–1,626. There were 21,307 parsimony-informative SNPs across all loci. Reducing our data matrix to evenly sampled populations, we retained 8,970 parsimony informative SNPs. We deposited aligned ddRADseq and UCE matrices on Dryad (doi: pending) and raw reads on NCBI’s Sequence Read Archive (SRR: pending).

### Phylogenetic Inference

Phylogenetic relationships were broadly concordant across different inference methods (**Figures 2, S8, S9**). Our maximum likelihood phylogeny, SVDquartets individual lineage tree, and SVDquartets species tree from ddRADseq SNP data supported the monophyly of the *E. difficilis* / *E. occidentalis* complex relative to *E. flavescens.* Both also recovered a fully-supported sister relationship between a clade containing all unadmixed individuals from from the Sierra Madre del Sur of Guerrero and Oaxaca in southern Mexico (hereafter “MXS”) and a clade containing individuals in the Transvolcanic Belt, Sierra Madre Occidental, and Sierra Madre Oriental of Mexico (“MXO”), individuals in the US Rocky Mountains and Great Basin (“USE”) and individuals in the US Sierra Nevada, Cascades, and Pacific Coast Ranges (“USW”).

**Figure 2:**
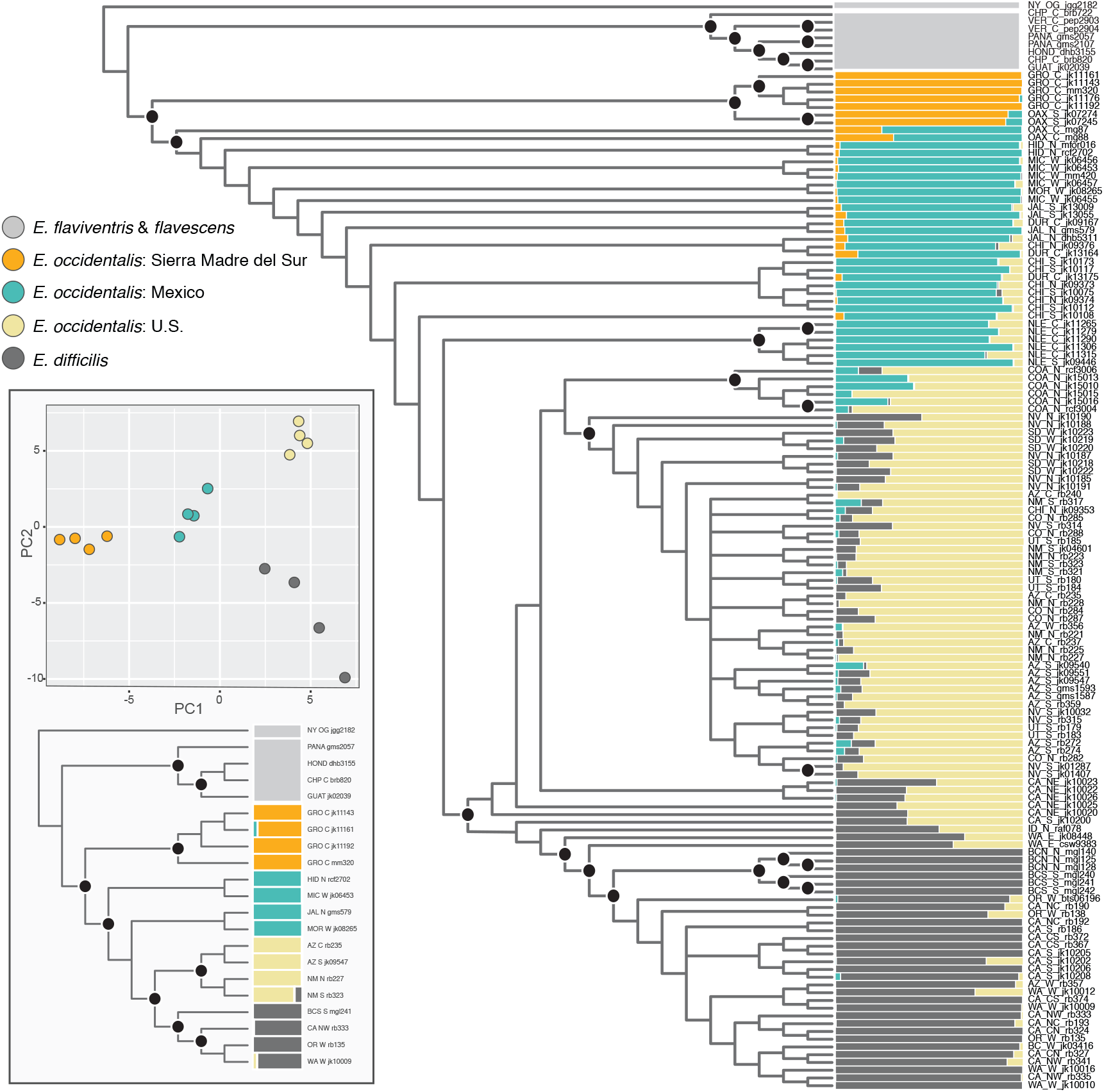
Population limits and phylogenetic relationships estimated using ddRADseq SNP data. (L): `SVDquartets` species tree of “pure” populations and PCA. (R): SVD quartets tree including all individuals and paired with `structure` ancestry proportions at *k*=4.

Within the *E. difficilis* / *E. occidentalis* clade, our species tree recovered fully supported subclades corresponding to USW and USE, nested within a grade of MXO individuals. Maximum likelihood inference recovered fully supported clades coinciding with the majority of USW, USE, and MXO individuals, but nested within a grade of individuals of putative hybrid ancestry (see below). Discordant nodes between maximum-likelihood and species trees lacked bootstrap support in all instances. A species tree inferred with our UCE SNP dataset recovered relationships within USE and MXO individuals concordant with those in our ddRADseq trees (**Figure S8**). Maximum likelihood inference with the mitochondrial gene ND2 differed only from SNP datasets by placing MXS individuals sister to a clade comprised of *E. flavescens* and USW / USE (**Figure S9**), rendering the latter paraphyletic. This placement, however, was not well supported.

### Population Structure Inference

Assessing clustering across multiple values of *k* with Bayesian Information Criteria revealed a single shift in slope at *k*=4. This result was corroborated by mean estimated likelihood values for assignment schemes in structure, which narrowly supported a four-population model over a three-population model (**Table 1**). Individual population assignments were broadly concordant between *k*-means and Bayesian clustering approaches. The primary difference was driven by `structure’s` ability to model “admixture” (e.g., to minimize assignment uncertainty by invoking proportional ancestry contributions from multiple populations), which identified a number of individuals of putative admixed ancestry. These admixed individuals were predominately found in northern Coahuila, Mexico, and northeastern California.

**Table 1:** Likelihood scores for different population assignment / species limit partitioning schemes (values of *k* in `structure`).

| $k$ | Replicates | Mean | Standard Deviation |
| --- | --- | --- | --- |
| 1 | 10 | -190860.17 | 2.23 |
| 2 | 10 | -183007.93 | 683.88 |
| 3 | 10 | -176774.65 | 34.33 |
| 4 | 10 | -176606.12 | 363.79 |
| 5 | 10 | -237514.34 | 111236.38 |
| 6 | 10 | -269824.40 | 129220.10 |

### Genomic Cline Analysis

For Transect 1, the best model included fixed scaling and exponential decay in the right end tail of the cline (AICc=-24.368). The best model for Transect 2 included fixed scaling and no exponential decay in either tail (AICc=-62.644). The best model for Transect 3 also included fixed scaling and no decay (AICc=-49.615). Transect 3 was narrowest, with a mean width of 73.789 km, followed by Transect 1, with a mean width of 210.078 km, and then Transect 2, with a mean width of 574.840 km (**Figure 4**). Full confidence intervals are provided in **Table S2**.

**Figure 3:**
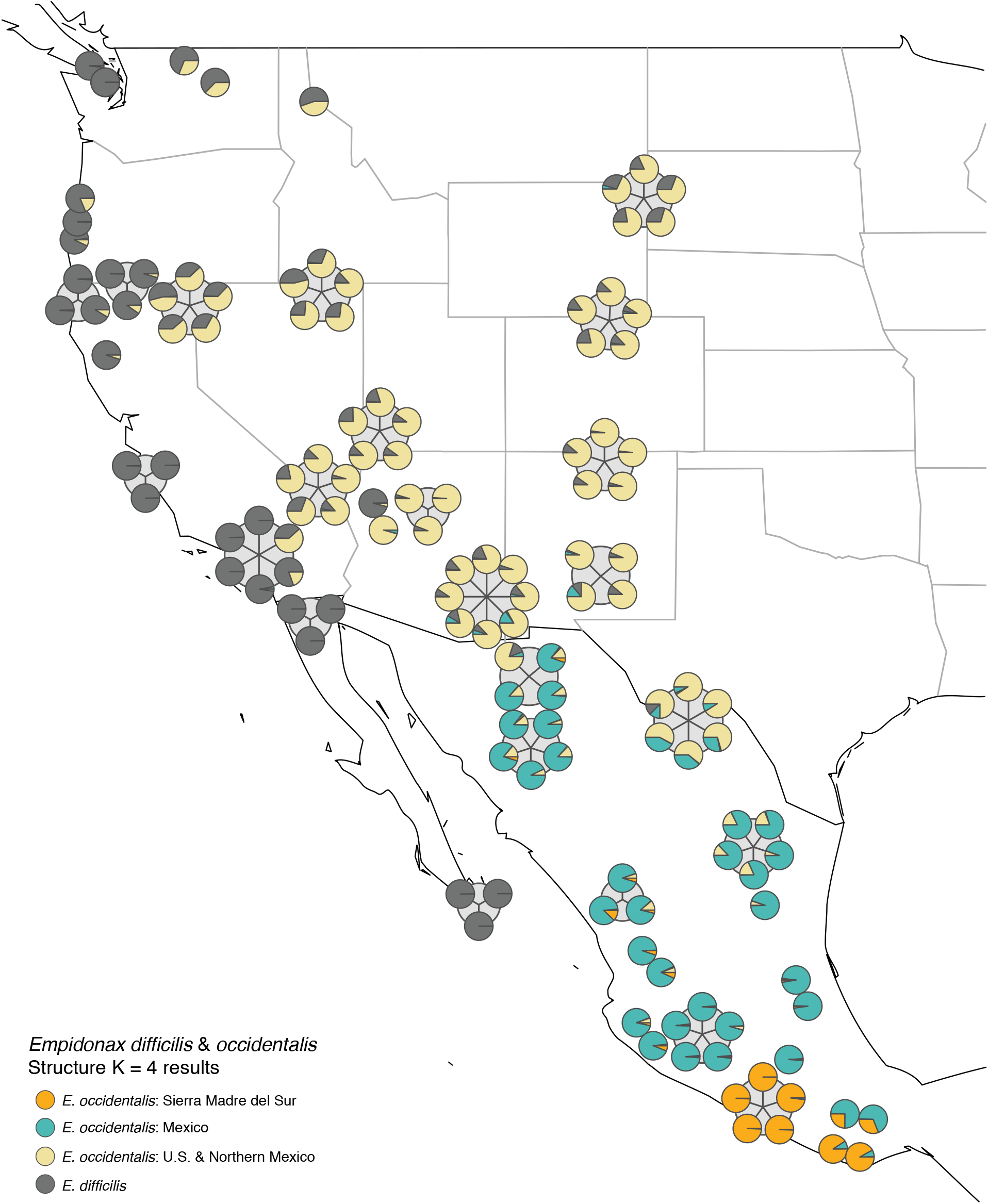
Proportional ancestry at *k*=4 for *Empidonax difficilis* and *E. occidentalis* individuals. Each circle represents an individual, with clusters of circles representing multiple individuals at a single sampling locality.

**Figure 4:**
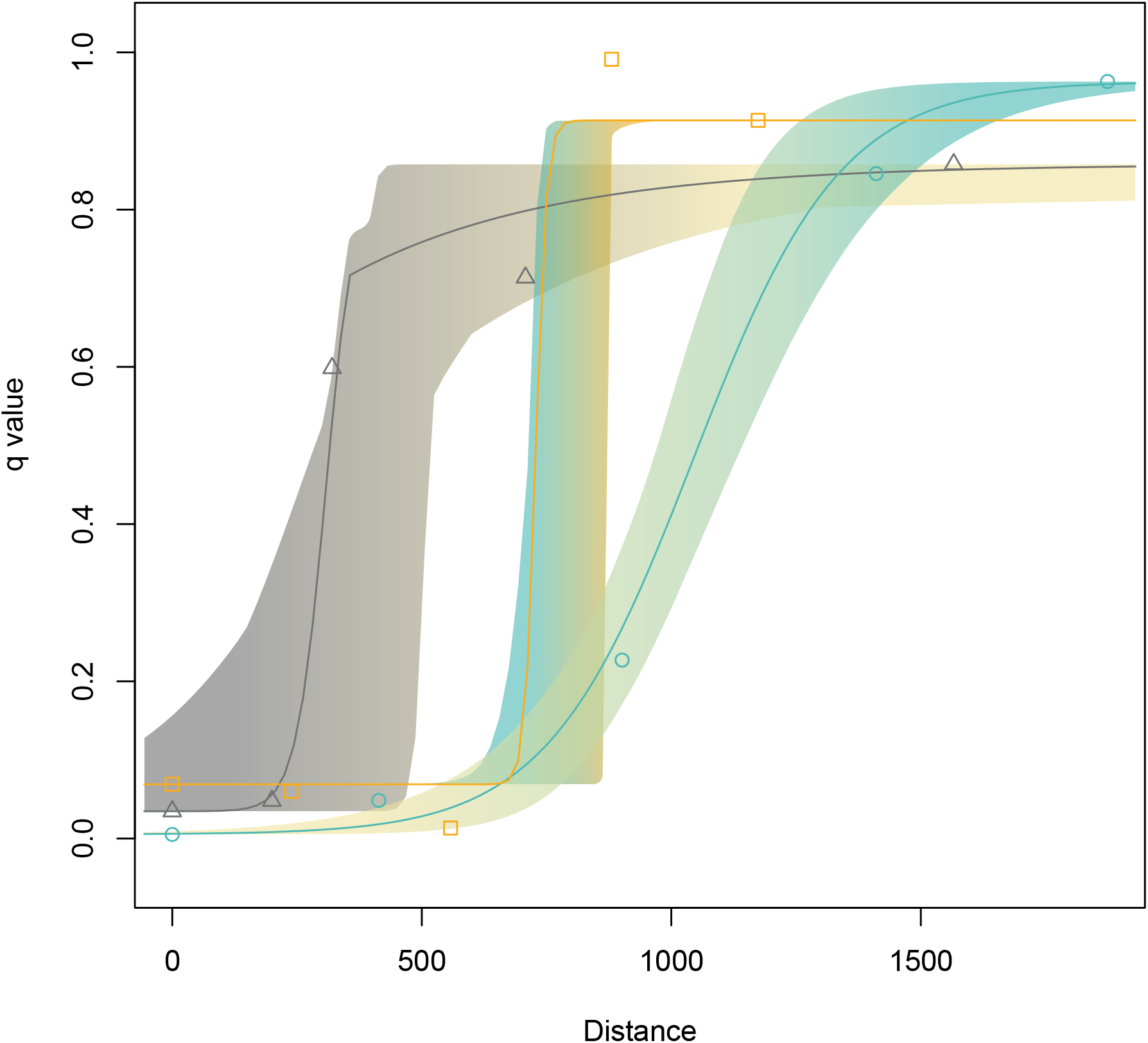
Genomic clines for transects spanning hybrid zones between *Empidonax difficilis* and *E. occidentalis* in NW USA (left, gray triangles; “Transect 1”); between Mexican *E. occidentalis* and resident *E. occidentalis populations* in the Sierra Madre del Sur (center, orange squares; “Transect 3”), and between US *E. occidentalis* and Mexican *E. occidentalis* (right, turquoise squares; “Transect 2”). *Q* value indicates admixture proportion from `structure` analyses.

## Discussion

### Geographic Bias in Sampling and Species Limits

Increased sampling of low-latitude populations revealed that the *E. difficilis* / *E. occidentalis* complex contains four parapatric lineages, all of which interbreed to varying degrees with neighboring populations and / or share a significant amount of ancestral polymorphism (**Figure 2**). The apparent paraphyly of *E. occidentalis* with respect to *E. difficilis* and previously unrecognized distinctiveness of the population of *E. occidentalis* in Mexico’s Sierra Madre del Sur provide an important cautionary tale for systematists and phylogeneticists attempting to proscribe species limits with incomplete geographic sampling. Few vertebrate faunas have been as intensively studied as North American birds, yet our results demonstrate that the presence of unsampled populations or sister species in less well-studied regions of a taxon’s range may dramatically alter species-delimitation schemes and inferred phylogenetic hypotheses—even when these unsampled lineages appear to be peripheral, small in population size, or otherwise ‘unimportant.’ The presence of strong latitudinal gradients in species richness (Willig et al. 2003) and intraspecific genetic diversity (Smith et al. 2017) increase the probability that many lower-latitude populations may represent distinct lineages. We encourage biologists working with taxa found across large latitudinal ranges to make efforts to include samples from the full extent of their distribution whenever possible, and consider the biases of missing samples when impossible.

Extremely limited phenotypic variation in tyrannid flycatchers broadly and *Empidonax* in particular has perplexed generations of naturalists, and raised questions about the degree and rate of evolution of reproductive isolation in birds (Zink and Johnson 1984). Our data suggest these studies have at least in part been hampered by a misunderstanding of the true evolutionary relationships and total diversity of these taxa. Would detailed study of other *Empidonax* alter our understanding of evolution across the genus, informing our understanding of trait evolution, reproductive isolation, and phylogenetic relationships? We encourage researchers to prioritize studies of this poorly known group to further resolve questions of species limits.

### Species limits, gene flow, and evolutionarily significant units

Our results highlight a conflict between phylogenetic and population genetic (clustering-based) species delimitation schemes that is likely to become increasingly important as genomic datasets reveal previously hidden histories of gene flow and incomplete reproductive isolation during population divergence (Zarza et al. 2016). A strictly phylogenetic species concept encounters two problems as applied to western *Empidonax* flycatchers that limit its utility. First, the species complex includes at least two regions of probable admixture among three geographically and genetically structured groups that appear strong enough to distort phylogenetic inference with both coalescent model-based and maximum likelihood approaches. While admixture proportions in `structure` analyses reflect uncertainty in assignment probability rather than explicitly modeling introgression, and may thus reflect the signature of incomplete lineage sorting rather than secondary contact and introgression in recent divergence events like those in *Empidonax*, we believe that the concentration of admixed individuals at zones of contact between geographically sorted genotypic clusters nonetheless implicates some degree of recent gene flow. Formal tests for introgression (Durand et al. 2011) or phylogenetic network analyses (Wen et al. 2018) could help evaluate the relative contributions of these sources of uncertainty. Second, *E. occidentalis* appears to be a paraphyletic taxon, taxon, composed of a grade in which pure and hybrid *E. difficilis* are nested (though low support for internal nodes may indicate biased inference from incomplete lineage sorting or reticulation; **Figure 2**). Nevertheless, support for a four taxa model from population genetic clustering shows that the division between northern and southern populations of *E. occidentalis* is not arbitrary: although not explicitly modeled in our study, these results suggest a reduction in gene flow between halves of the species range by some combination of habitat limitations, geographic isolation, or incipient reproductive isolation. Regardless of the epistemic reality of species themselves, speciation is clearly a dynamic and continuous process, and evolutionarily significant units do not need to adhere to a strictly bifurcating tree (Turgeon and Bernatchez 2003).

Across *Empidonax* more broadly, cline analysis refutes suggestions of a linear relationship between phenotypic divergence and reproductive isolation (Johnson 1980). MXS populations are essentially indistinguishable from adjacent MXO populations with many standard morphometric measurements (Johnson 1980; Klicka et al. unpublished), but show a steep transition in allele frequencies (**Figure 4**) indicating limited gene flow between groups relative to the more morphologically divergent but less isolated pair of USE and USW. Our study shows the *E. difficilis* / *E. occidentalis* complex continues to illustrate this “gray area” of contemporary speciation research and species theories by suggesting several confounding processes may be at play: reticulation of lineages, local adaptation, differential rates of trait evolution, and unobserved behavioral differences.

*Evolution and Biogeography of* Empidonax difficilis *and* E. occidentalis: The range of western North American *Empidonax* flycatchers is typical of an array of roughly codistributed taxa restricted to mountainous Nearctic habitats characterized by forests of pine, oak, and fir. Because much of northern North America was glaciated during the Pleistocene, phylogeographic patterns in these species have been attributed to the expansion out of multiple, isolated, ice-free refugia (Klicka and Zink 1999). Though likely locally distorted by the effects of gene flow, our phylogeny of western *Empidonax* is at least geographically consistent with this model: with Central American *E. flavescens* and *E. flaviventris* as sister taxa, we see sequential splits to the southernmost portion of the range of E. occidentalis in the Sierra Madre del Sur, a split between the Sierra Madre del Sur and the Sierra Madres Occidental / Oriental, the Transvolcanic Belt, and the U.S. Rocky Mountains, followed by a split between these ranges and Pacific Coast Ranges, with continued interbreeding at contact zones of suitable habitat. We believe this sequence most likely reflects dispersal and colonization followed by allopatric divergence, although vicariance and secondary contact through the contraction of appropriate montane habitat as a result of climatic cycling is at least possible. Regardless of the mode of divergence, these data are consistent with previous studies of diversification across the Mexican highlands (Bryson et al. 2010) and with the conclusion that the Sierra Madre del Sur is a hotspot of phylogenetic endemism (Blancas-Calva et al. 2010; van Els et al. 2014), and we encourage increased attention to this distinctive region.

Are these lineages evolving neutrally, or adapting to the broadly varied climate space across the group’s range? Satisfactorily answering this question is beyond the scope of the current paper, but we wish to emphasize the role of topography in isolating *E. difficilis* / *E. occidentalis* populations across their range: each of the four clusters identified in our analysis broadly coincides with a complex of mountain ranges (the Cascades / Sierra Nevada / Coast Ranges, the Rocky Mountains, the Sierra Madre Occidental / Oriental, and the Sierra Madre del Sur) separated by less suitable lowland habitat (the Great Basin, the Sonoran and Chihuahuan Desert, and the Balsas Basin) though linked by discontinuous ‘sky islands’ in some places. These barriers have been implicated in restricting gene flow in other terrestrial vertebrate taxa (Bryson et al. 2010; Gowen et al. 2014; van Els et al. 2014; Mastretta-Yanes 2015). In light of previous research suggesting ecologic diversification is responsible for divergence of *Empidonax* north of the Mexican border (Johnson and Cicero 2002), however, local adaptation may be playing a role in further reducing gene flow across these barriers.

### Taxonomic Recommendations

Our results conflict with contemporary taxonomic treatments, and are consistent with a range of one to four species depending on the invoked species concept. As discussed above, a strict phylogenetic species concept is difficult to apply in the face of probable reticulation, but identifies either a single, widely distributed species (if putative hybrid individuals are included) or two species (if hybrids are excluded), with one lineage inhabiting the Sierra Madre del Sur and the other the remainder of continental North America. A genotypic clustering species definition (Mallet 1995) identifies four distinct species, coinciding with our USW, USE, MXO, and MXS regional definitions. This assignment scheme is further supported by genomic cline analysis: while cline width varies across our sampled transects, either as a consequence of different levels of reproductive isolation or the availability of samples, the underlying sigmoidal model for all three clines is consistent with some degree of selection against hybrids (Endler 1977). We recognize that a pragmatic approach to species delimitation for end users (e.g., field naturalists, conservationists, and ecologists) needs to incorporate diagnostic phenotypic characters, whether in range, ecology, behavior, song or morphology. A detailed analysis of these data will be presented elsewhere.

### Conclusions

Re-examining a classic morphologically cryptic sibling species complex with genome-wide data and expanded geographic sampling, we identify a highly distinct unrecognized lineage in a previously unsampled portion of the group’s range. We additionally identify the paraphyly of a widely recognized species, although this conflicts with the best-supported species-delimitation model. Our phylogeny supports a model of expansion from southern Mexico. Our results highlight the pervasive impacts of biased geographic sampling, even in well-studied vertebrate groups like birds, and illustrate what is a common problem to phylogenetic species concepts in the face of recent divergence or reticulate evolution.

### Data availability and Supplemental Material

Supplemental material, code, and sequence alignments are available via Dryad (pending). Raw sequence data are available from the NCBI SRA (pending).

## Acknowledgements

We thank the Burke Museum of Natural History and Culture, the American Museum of Natural History, the Universidad Nacional Autónoma de México, and the Denver Museum of Nature and Science for providing tissue samples. E.B.L. was supported by a NDSEG Fellowship from the U.S. Department of Defense.

## Supplemental Material

Tables S1 and S2 available via Dryad (link pending)

**Figure 5:**
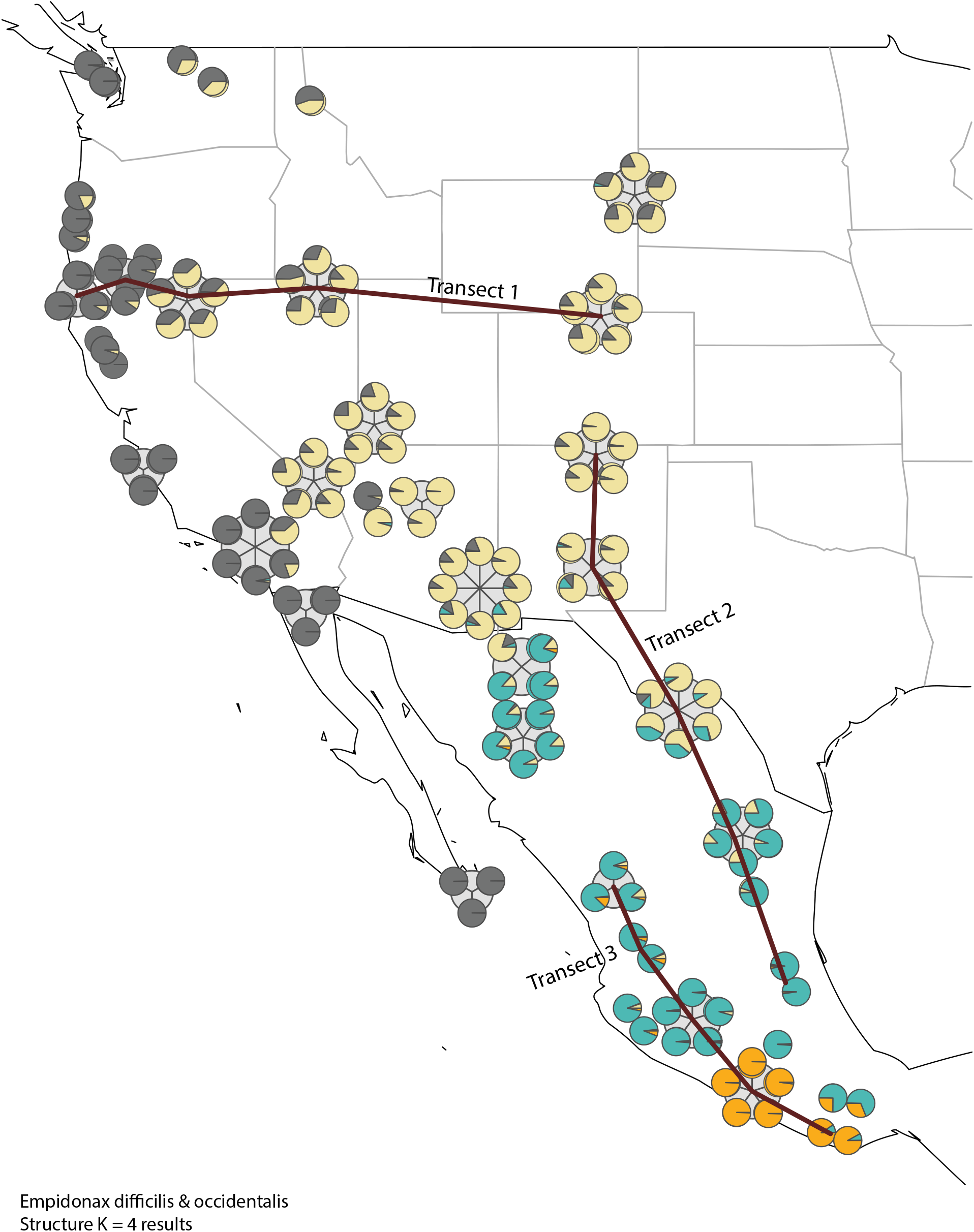
Transects for genomic cline analysis.

**Figure 6:**
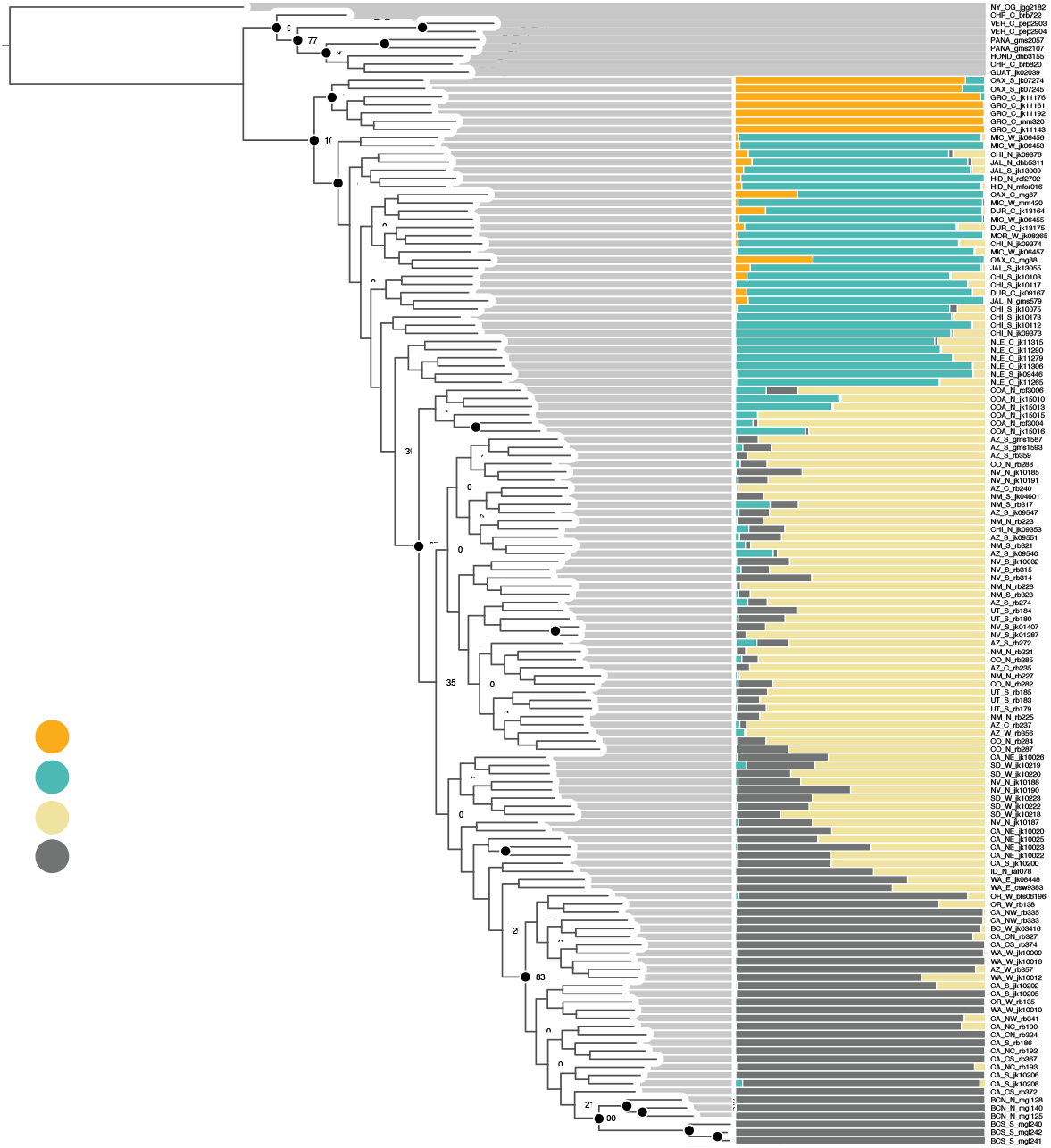
`RAxML` phylogeny from concatenated RAD loci and `structure` admixture proportions from RADseq SNP data.

**Figure 7:**
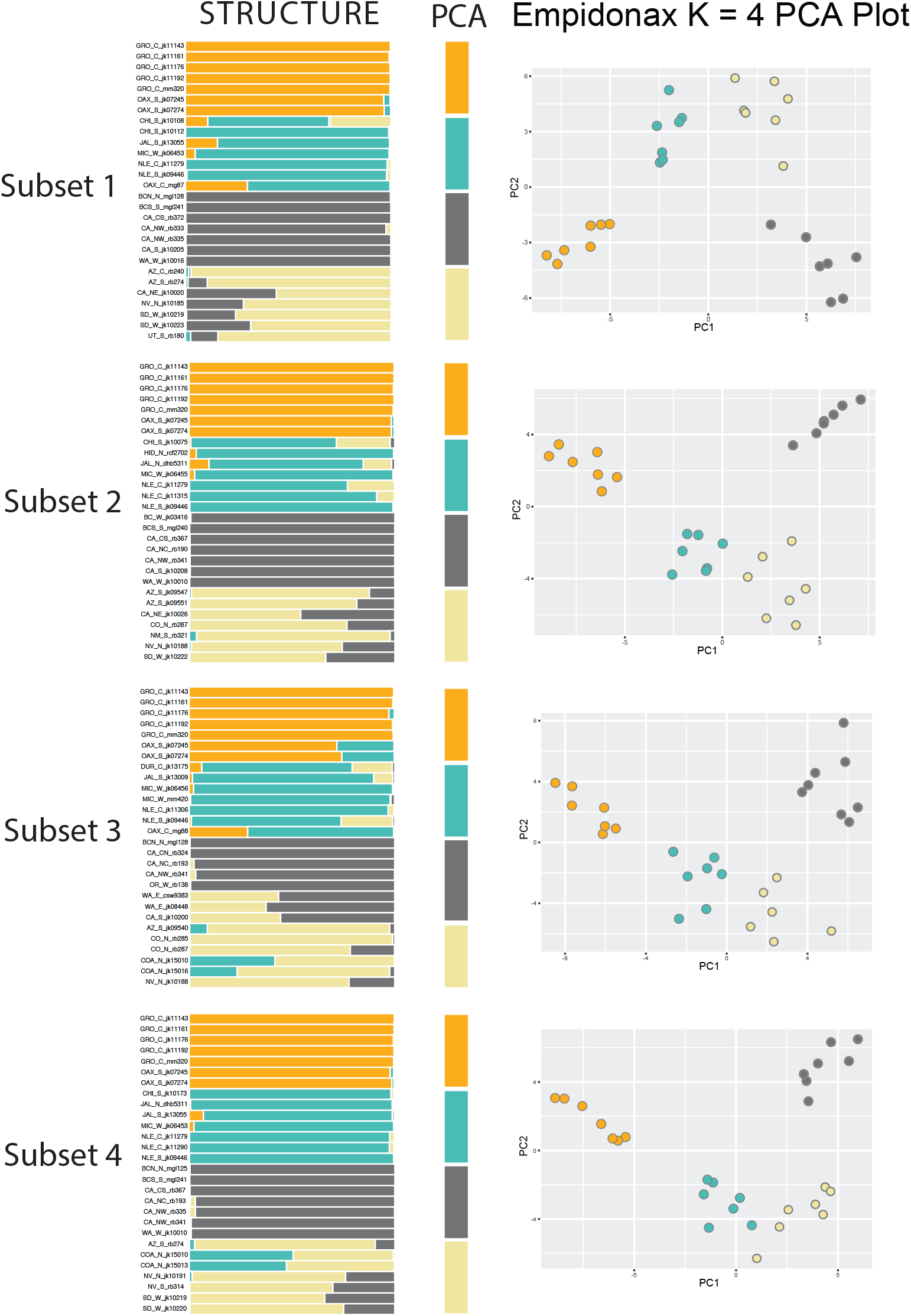
Population structure inference results from equal sample sized subsets, RADseq SNP data.

**Figure 8:**
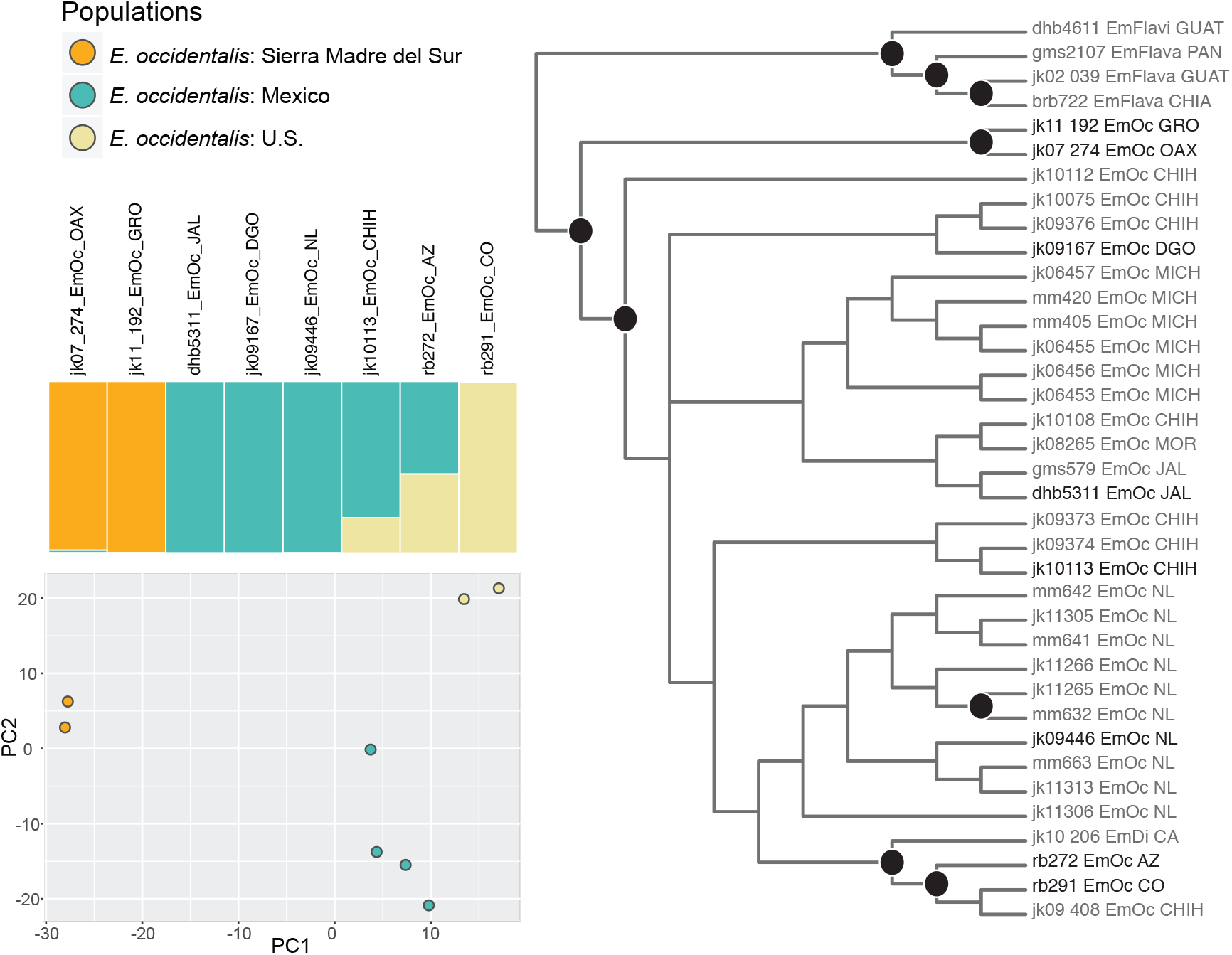
UCE-based `SVDquartets` phylogeny and PCA

**Figure 9:**
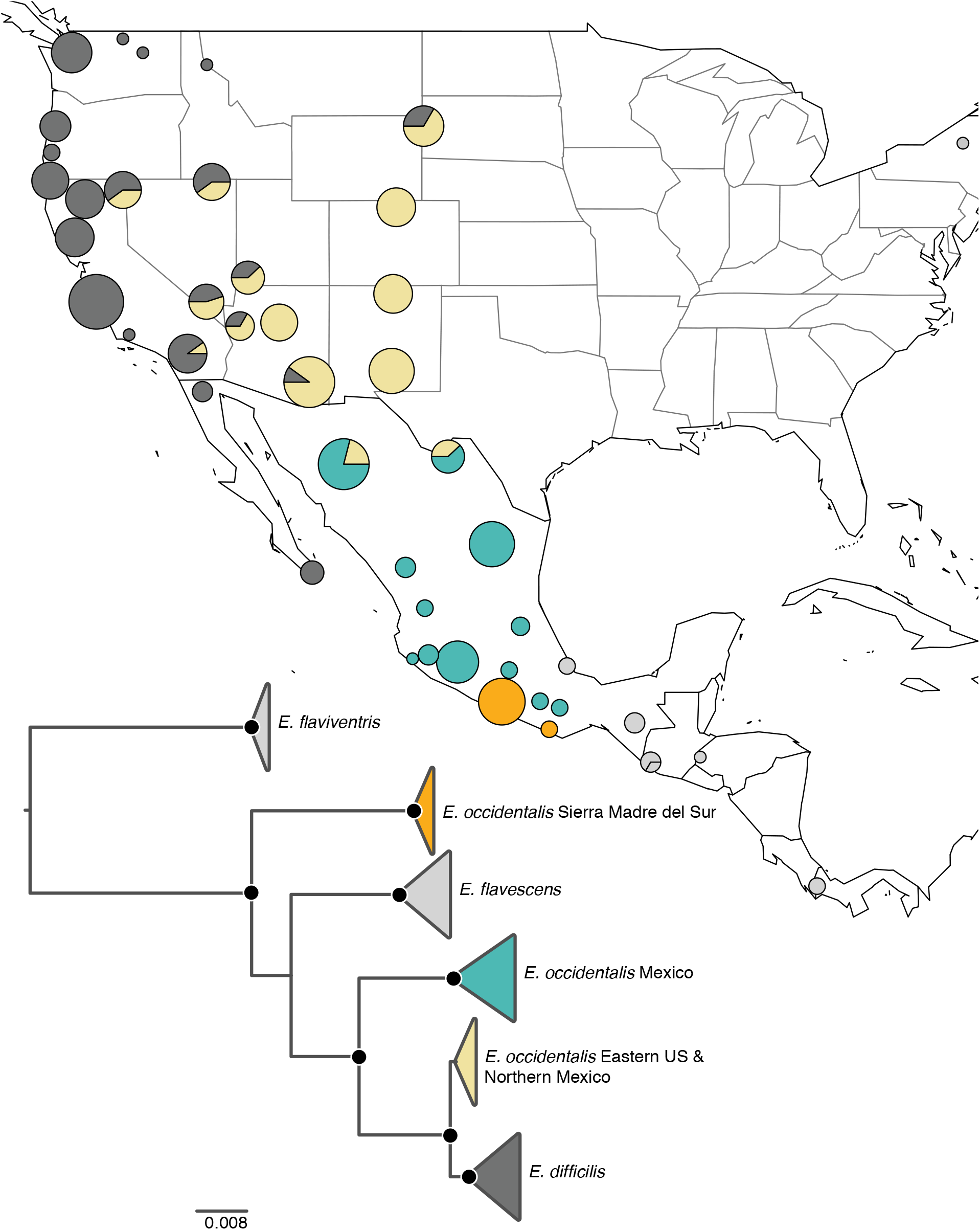
ND2-based `RAxML` phylogeny

